# Structural basis of promiscuous inhibition of *Listeria* virulence activator PrfA by oligopeptides

**DOI:** 10.1101/2024.05.24.595734

**Authors:** Tobias Hainzl, Mariela Scortti, Cecilia Lindgren, Christin Grundström, Emilia Krypotou, José A. Vázquez-Boland, A. Elisabeth Sauer-Eriksson

## Abstract

The facultative pathogen *Listeria monocytogenes* uses a master regulator, PrfA, to tightly control the fitness-costly expression of its virulence factors. We found that PrfA activity is repressed via competitive occupancy of the binding site for the PrfA-activating cofactor glutathione by exogenous nutritional oligopeptides. The inhibitory peptides show different sequence and physicochemical properties, but how such wide variety of oligopeptides can bind PrfA was unclear. Using crystal structure analysis of PrfA complexed with inhibitory tri- and tetrapeptides, we show here that the binding promiscuity is due to the ability of PrfA β5 in the glutathione-binding tunnel to establish parallel or antiparallel β-sheet-like interactions with the peptide backbone. Spacious tunnel pockets provide additional flexibility for unspecific peptide accommodation while providing selectivity for hydrophobic residues. Hydrophobic contributions from two adjacent peptide residues appears to be critical for PrfA inhibitory binding. In contrast to glutathione, peptide binding prevents the conformational change required for PrfA activation and formation of the DNA-binding helix-turn-helix motifs, effectively inhibiting virulence expression.

## Introduction

*Listeria monocytogenes*, the causative bacterium of foodborne listeriosis, is an archetypal facultative pathogen that can live freely in the environment or parasitically in mammalian cells^1-3^. Maintaining optimal fitness across this dual lifestyle is essential for the competitiveness and evolutionary viability of this type of pathogens. This is largely achieved by master regulators that turn virulence genes on during infection and switch their expression off outside the host^4-6^. In *Listeria*, this role is fulfilled by the master regulator PrfA, a bacterial Crp/Fnr transcription factor^7-10^. PrfA controls ten key virulence genes, the so-called PrfA regulon^11,12^, which are strongly induced intracellularly and repressed during saprophytic growth^5,13-15^.

PrfA-dependent expression is modulated by signals thought to allow *L. monocytogenes* to sense the saprophyte to pathogen transition^13,14^. For example, an RNA thermoswitch inhibits *prfA* gene translation at temperatures below 30°C, as would be found outside a warm-blooded host^16^. PrfA-regulated genes are also repressed upon utilization of certain sugars, in particular plant-derived β-glucosides such as cellobiose^17^, presumably abundant in the soil habitat of *L. monocytogenes*^18^. Free fatty acids also interfere with the DNA-binding activity of PrfA^19^. In contrast, virulence gene expression is upregulated by other cues. These include a reducing environment as well as stress signals or poor amino acid availability via the SigB and CodY regulators^20-22^. To what extent the above PrfA-modulating mechanisms contribute to the strong activation of listerial virulence genes observed in host cells remains uncharacterized.

A major step in understanding the listerial virulence gene “on-off” switching was the discovery that PrfA activity *in vivo* requires the redox tripeptide glutathione (γ-L-glutamyl-L-cysteinylglycine, GSH)^23^. In *L. monocytogenes*, GSH is endogenously synthesized by the glutathione synthase, GshF^24^, and an intact *gshF* gene in addition to PrfA is essential for virulence^23,25^. PrfA is a dimeric protein of 54.5 kDa formed by two identical monomers, each comprising two distinct domains connected via an interfacial α-helix linker^26,27^. The N-terminal domain is an eight-stranded cyclic nucleotide binding domain (CNBD) β-sandwich, of unknown function in PrfA. The C-terminal domain contains the DNA-binding helix-turn-helix (HTH) motif. Each of the two HTHs binds to one of the arms of the 14-bp palindromic sequence called the “PrfA-box” located at the -35 region of the target promoters^8,12,28^. GSH binds to PrfA with low affinity (K_*d*_ ≈4 mM)^23^, in a large tunnel between the N- and C-terminal domains^29^. GSH binding to each PrfA monomer stabilizes the dimer in the active “on” conformation with the HTH motifs correctly folded for productive interaction with the PrfA-box^29^.

While these studies identified GSH as a critical PrfA-activating co-factor, it was still unclear how GSH-dependent PrfA activation is controlled. We recently elucidated that the PrfA-GSH system is regulated by the composition of oligopeptides imported from the surrounding medium via the listerial Opp permease^25^, as follows: peptides that provide the essential GSH precursor cysteine activate PrfA; conversely, peptides lacking cysteines directly inhibit PrfA. Through this mechanism, *L. monocytogenes* exploits oligopeptides—an abundant and critical nitrogen source for microbial growth—to sense its habitat and control virulence gene expression by balancing their antagonistic effects on PrfA activity^25^.

Co-crystallization with an example inhibitory peptide, leucylleucine (LL), revealed that the dipeptide bound in the GSH-binding site in one of the PrfA monomers and had the same extended conformation as the GSH tripeptide. However, LL binding prevented the correct positioning of PrfA C-terminal DNA-binding helices^25^. Interestingly, tested inhibitory peptides ranged from di- to octapeptides with chemical properties including both negatively and positively charged residues as well as hydrophobic, polar, and aromatic residues. To gain detailed insight into the molecular mechanisms of peptide-mediated inhibition of PrfA, we in this study determined and analysed the crystal structures of sequence-diverse peptides in complex with PrfA. We found that the peptides bound at the GSH-binding site, also referred to as the tunnel site, by forming parallel or antiparallel main-chain main-chain interactions with the CNBD β-strand β5. In addition, hydrophobic pockets at the tunnel site provide some degree of selectivity for binding by interacting with hydrophobic or aromatic side chains of two consecutive peptide residues. Our findings provide structural insights into the unique ability of PrfA to accommodate a wide range of peptides with structurally and chemically diverse amino acid side chains.

## Results

### PrfA-peptide co-crystallisation and biophysical analyses

We succeeded in determining the crystal structures of PrfA in complex with seven peptides of various chemical properties^25^ (Fig. 1). For data collection and refinement statistics see Supplementary Table 1. Weak electron density for bound ligands due to conformational flexibility or low occupancy is a common problem in protein-ligand crystal structures. The use of bulk-solvent models in crystallographic refinement may also obscure densities in areas not occupied by protein atoms^30^. To overcome these limitations, in this study we used polder maps^30^ in addition to classical difference maps. The LigandFit step in the procedure provided local correlation coefficients (CC) values over 0.75, which support binding of the peptide ligands to PrfA in all the complexes studied (Supplementary Table 2). Fig. 2a illustrates the improvement of interpretability with the polder map of the tripeptide LLL as an example. Polder maps for the remaining peptides are shown in Supplementary Fig. 1.

**Fig. 1.**
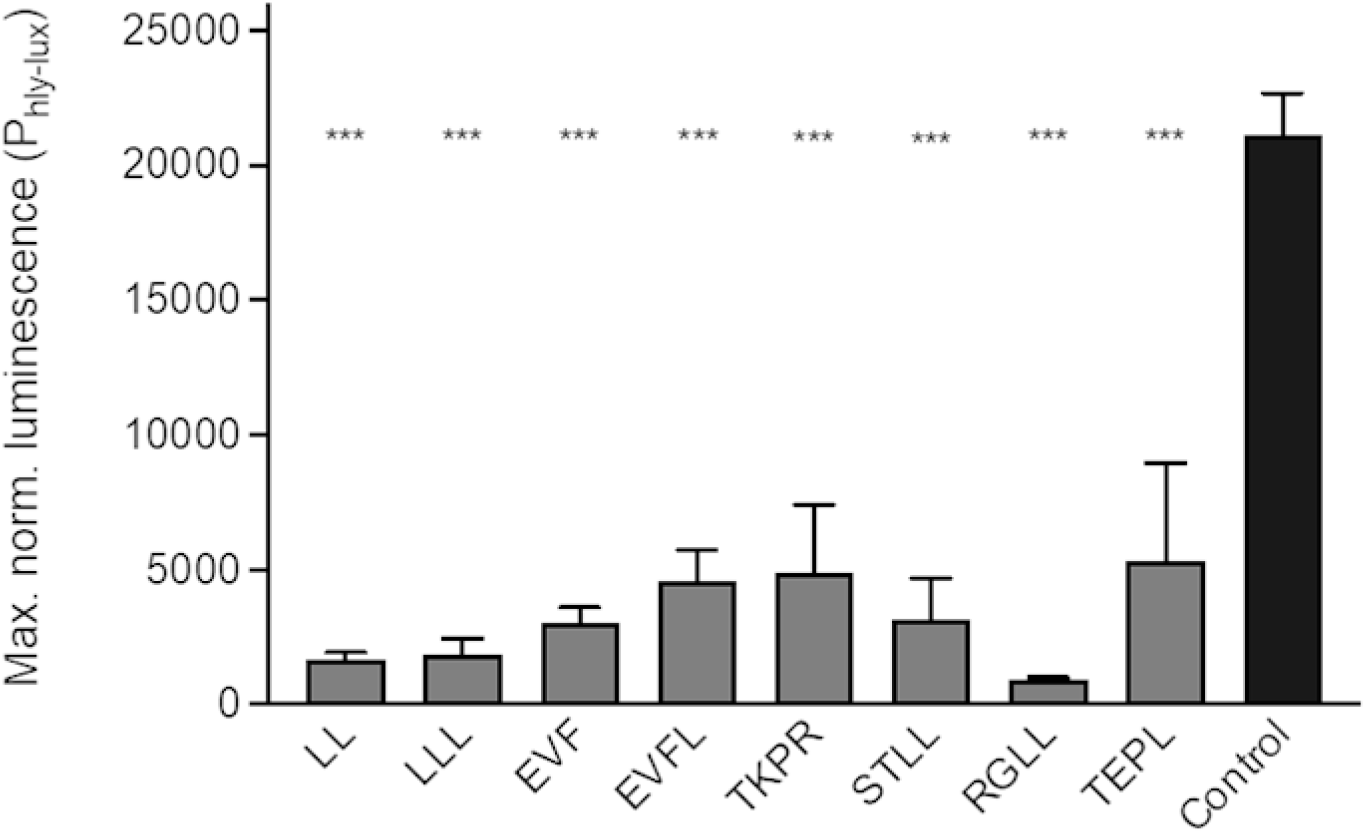
PrfA inhibitory activity of peptides. PrfA-dependent virulence gene expression of *L. monocytogenes* P14-Phly-lux quantified as maximum luminescence normalized to bacterial growth (OD600). Peptides were added at 1 mM concentration. Inhibitory peptides LL, LLL, EVF, EVFL and TKPR were previously described^28^. Means ± SEM from at least three experiments in triplicate. Asterisks indicate p < 0.001 relative to control (medium without peptides). One-way ANOVA and Dunnett test for multiple comparisons.

**Fig. 2.**
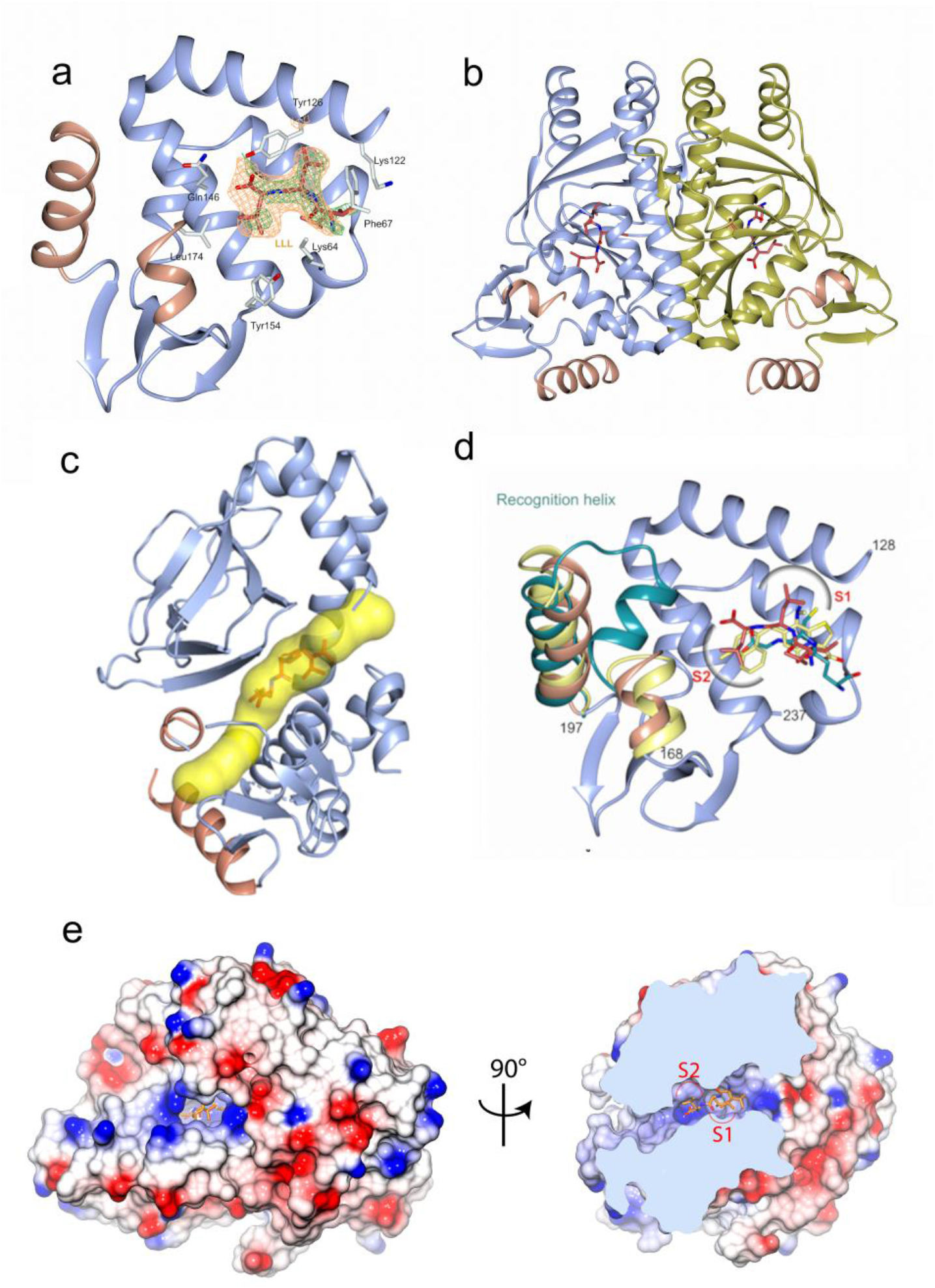
Binding of oligopeptide LLL to the interdomain tunnel of PrfA. (**a**) Ribbon representation of the C-terminal domain of monomer A of the PrfA-LLL complex. Residues 128-237 are shown in blue with the HTH motif (residues Asn168-Lys196) highlighted in orange. For clarity, only a few selected residues are shown as sticks. The LLL peptide is shown in crimson red. The first leucine residue of the peptide has two conformations. The difference (|Fo|-|Fc|) and polder electron density maps are green and orange, respectively. The maps are contoured at three times the RMSD value of the map, covering the LLL peptide only. (**b**) Ribbon representation of the PrfA homodimer in complex with LLL. Monomers A and B are shown in blue and gold, respectively, the HTH motif is highlighted in orange, and the LLL peptide is crimson red. (**c**) Structure of the PrfA-LLL complex showing a PrfA monomer in cartoon representation with the main tunnel (calculated with CAVER^69^) as yellow surface. (**d**) The figure outlines the position of the hydrophobic pocket S1 and S2 at the tunnel site of the PrfA-LLL complex. For clarity no protein residues are shown. Included in the figure are superpositioned structures of PrfA bound to LL (yellow, PDB code 6hck^28^) and PrfA bound to the activator molecule GSH (dark cyan, PDB code 5lrr^27^). (**e**) Two orientations of an electrostatic surface representation of the PrfA dimer with LLL bound at the tunnel site of monomer A.

To gain functional insight into the direct interaction between PrfA and the peptides, the binding thermodynamics was investigated using isothermal titration calorimetry (ITC). The results, shown in Supplementary Fig. 2, indicated that the peptides bind to PrfA with a negative enthalpy change. Analysis of the integrated heat peaks, as a function of ligand-to-protein ratio, showed that peptides LLL and LL have the strongest affinity, with dissociation constants (K_d_) in the low μM range. Following these, peptides EVF and EVFL have K_d_ values of 12 μM and 15 μM, respectively, while peptides STLL and RGLL bind significantly weaker with K_d_ values of 32 μM and 145 μM, respectively. The binding affinities of peptides TKPR and TEPL could not be reliably quantified under the experimental conditions used. GSH binding was undetectable by ITC, as previously reported^25^, probably owing to its weak affinity to PrfA^23^.

To analyze how peptide binding affects the DNA-binding properties of PrfA, we performed biolayer interferometry (BLI) experiments. All seven peptides inhibited DNA binding. Consistent with the ITC data, peptides LLL, LL, and EVF were found to be the strongest inhibitors, with peptides LLL and LL completely blocking DNA binding, as shown in Fig. 3. Peptides AG and LIVA which showed no significant PrfA-inhibitory activity *in vivo* in *L. monocytogenes* were used as controls. Additional BLI experiments, performed both with and without the activator molecule glutathione (GSH), showed that GSH can displace bound oligopeptides, confirming a competitive interaction (Fig. 3).

**Fig. 3.**
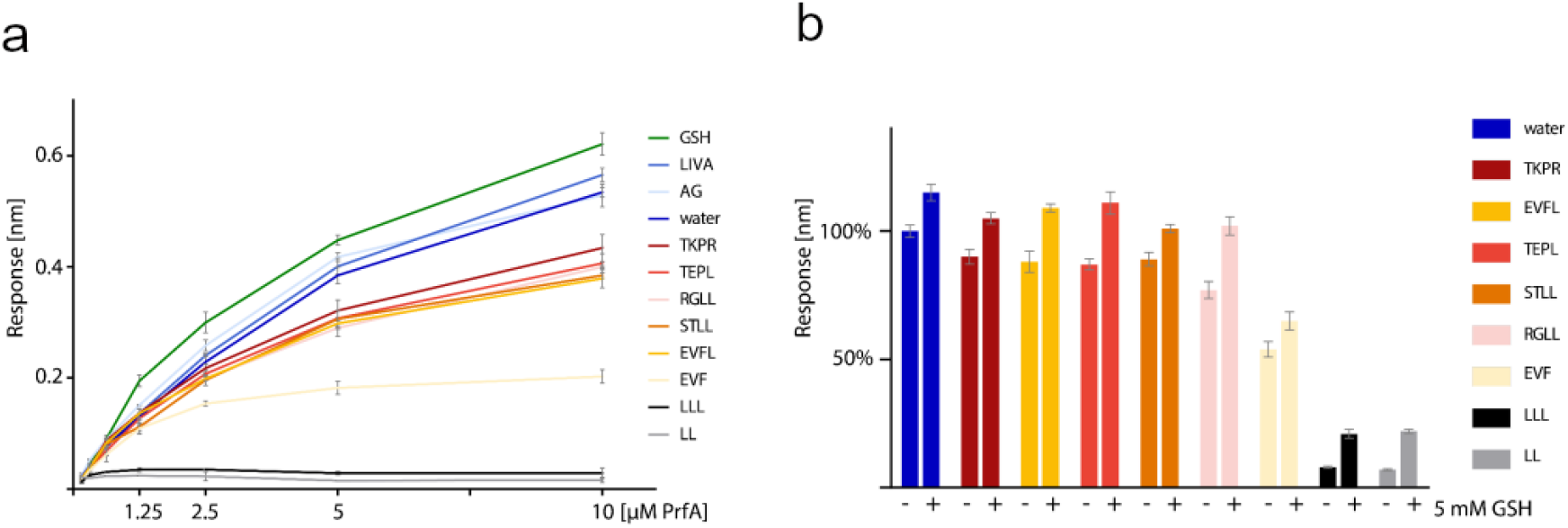
Modulation of PrfA-DNA binding by oligopeptides and GSH. (A) Inhibition of PrfA-DNA binding by oligopeptides. Effect of inhibitory and non-inhibitory oligopeptides on the binding of PrfA to PplcA/hly PrfA box DNA, measured by biolayer interferometry with a PrfA:oligopeptide ratio of 1:100. For GSH, a concentration of 5 mM was maintained for all PrfA concentrations. The graphs show the binding response plotted against PrfA concentration, with mean values ± standard deviation (SD). (B) Competitive modulation by GSH. The graph shows the enhancement of DNA binding of PrfA-oligopeptide mixtures (5 μM PrfA-500 μM oligopeptides) in the presence of 5 mM GSH. The Y-axis represents the binding response, with the mean response of PrfA alone set to 100%. Mean values ± SD are shown.

### Peptide binding at PrfA’s interdomain tunnel

PrfA is a 237-residue protein in which residues 1-109 and 138-237 constitute the N- and C-terminal domains, respectively, connected by the linker-helix spanning residues 110-137. Our previous studies showed that the binding site for the GSH cofactor^29^ and the dipeptide LL^25^ is positioned at the so-called interdomain tunnel or tunnel formed between the N- and C-terminal domains of the PrfA monomers. Although not all amino acids of the different peptides could be modelled (discussed below), all peptides in this study bound at this site. As an example, Fig. 2b and c show the structure of the PrfA-LLL complex with the tripeptide bound to the tunnel site of monomers A and B. Likewise, all other peptides bound to the tunnel site in monomers A and B with one exception TEPL (Supplementary Fig. 3). This tetrapeptide bound only to monomer A as did the previously studied LL dipeptide^25^. This suggests structural negative cooperativity in the binding of peptides to the PrfA dimer, where binding of one ligand in one monomer affects the shape of the other monomer in such a way that ligand binding affinity decreases, as observed in previous studies of PrfA in complex with 2-pyridone PrfA inhibitor^31,32^.

In the previous structure-guided 2-pyridone inhibitor studies, two binding pockets at the tunnel site, S1 and S2, were predicted to allow for some degree of selectivity in the PrfA-ligand interaction^32^. Selectivity pocket S1 creates possibilities for hydrophobic interactions with the side chains of PrfA residues Tyr63, Phe67, Tyr126, and Trp224 whereas pocket S2 does the same for Ile45, Tyr62, Ile149, Leu150, Tyr154, and Leu174. All PrfA-inhibitory peptides studied here had side chains that bound to the S1 and S2 pockets. The same has been previously reported for the binding of the dipeptide LL to PrfA^25^, confirming the functional importance of the S1 and S2 pockets in the interdomain tunnel for PrfA-peptide interaction (Fig. 2d-e and Supplementary Fig. 3).

### Interaction in parallel or anti-parallel β-sheet-like conformations underpins peptide binding flexibility

Except for TKPR, all peptides in this study are in an extended β-strand-like conformation: they establish main-chain contacts with PrfA strand β5 (Gln61-Lys64) and the turn leading to β6 (Gly65-Phe67) (Supplementary Fig. 3). A close-up view of β5 shows that it can form five main-chain hydrogen bonds with an incoming peptide (Fig. 4a). In addition, due to the cis-peptide bond between residues Gly65-Ala66, there are possibilities for hydrogen bonding with the main-chain carbonyl oxygen of Ala66. Five of the peptides—EVF, EVFL, LLL, STLL and TEPL—bind antiparallel to β5 by forming five main-chain hydrogen bonds (Fig. 4b, Supplementary Fig. 3). The polder maps showed that the first residue in both STLL and TEPL is flexible and could not be modelled; thus the modelled β5-binding residues for these peptides are –TLL and –EPL, respectively. EVFL, which is a one-amino acid extension of EVF, binds its first three residues identically to those of the tripeptide. In addition, the main chain atoms of Leu4 in EVFL make hydrogen bonds to the side chains of Gln146 and Tyr126, and its side chain packs against that of Leu174.

**Fig. 4.**
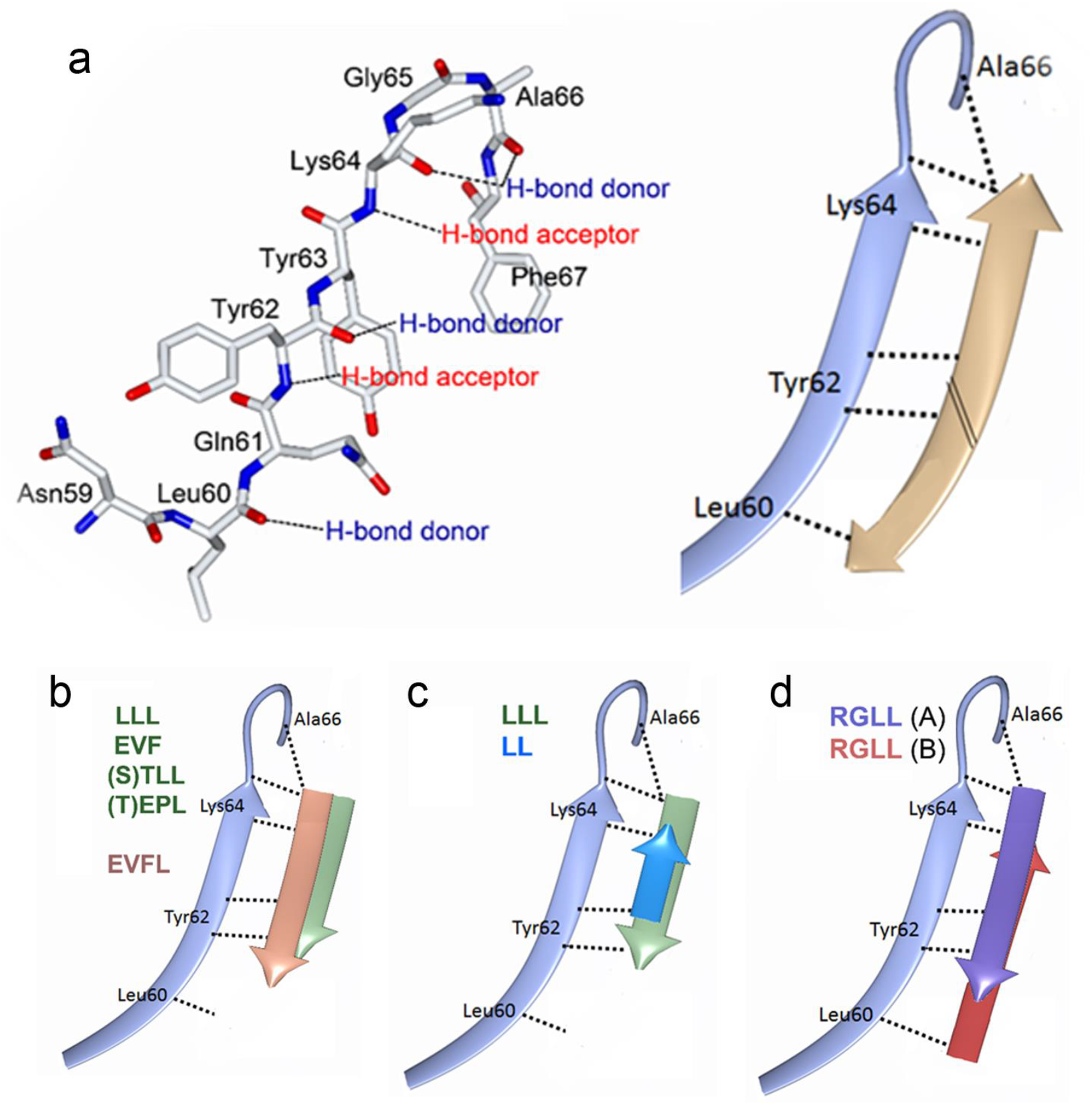
Peptide binding variability in PrfA. (**a**) Left: Drawing of beta strand β5 in PrfA showing six hydrogen-bonding possibilities to an incoming peptide. Right: These bonds can be used for both parallel and anti-parallel β-sheet formation with an incoming peptide. (**b-d**) Ribbon drawings showing how different peptides in this study bind parallel or anti-parallel to β5. The RGLL peptide binds anti-parallel in monomer A and parallel in monomer B.

Interestingly, while the Leu dipeptide bound parallel to β5 in a previous PrfA-LL crystal structure (PDB code 6hck^25^), the Leu tripeptide binds in an anti-parallel conformation in our PrfA-LLL complex (Fig. 4c). The side chains of the second and third leucines in the tripeptide bind identically to the Leu side chains of the dipeptide at the S1 and S2 sites. Both peptides form two hydrogen bonds each to the main chain carbonyl oxygen and amide nitrogen atoms of Tyr62 and Lys64, respectively (Supplementary Fig. 3). In addition, the N-terminal nitrogen atom of LL forms a hydrogen bond to the hydroxyl group of Tyr126. The C-terminal carboxyl group of LLL, in turn, forms hydrogen bonds to the main chain nitrogen atom and the Ne2 atom of Gln61. The only obvious structural difference between how LL and LLL bind to PrfA is the location of Gln146 positioned in the vicinity of the HTH motif; it is closer to the peptide in the PrfA-LL complex (Supplementary Fig. 3).

Another peptide that binds parallel to β5 is RGLL. Strikingly, this peptide binds in both directions within the same dimer—anti-parallel to β5 in monomer A and parallel to β5 in monomer B (Fig. 4d). However, in both monomers only the last two leucine residues in the tetrapeptide are well defined in the electron density (Supplementary Fig. 1). In the anti-parallel arrangement, the hydrogen-bonding pattern to β5 is similar to the ones observed in the tripeptides, e.g. LLL and EVF. However, the C-terminal carboxyl group does not make a hydrogen bond to the main chain nitrogen atom of Tyr62; rather it forms hydrogen bonds to the side chains of Gln61, Tyr126, and Gln146 (Supplementary Fig. 3). In monomer B, parallel binding of the peptide is made possible by hydrogen bonding to the main-chain oxygen atom of Gln61, the nitrogen and oxygen atoms of Tyr62, and the nitrogen atom of Lys64. In addition, there are interactions between the carbonyl oxygen of Arg1 of the peptide and the side chains of Lys130, and between the guanidinium group of Arg1 and the hydroxyl group of Ser142 (Supplementary Fig. 3).

Thus, the two binding modes of the LL and LLL peptides, and in particular the tetrapeptide RGLL in the same dimer, highlight that slightly alternating hydrogen bonding to the PrfA protein accommodates similar peptides or even the same one’s in two different conformations.

### Hydrophobic motifs at PrfA’s tunnel site contribute to inhibitory peptide binding

Superimposition of monomer A of the PrfA-LL and PrfA-LLL complexes shows that binding, in addition to main-chain β-strand formation, also involves interactions between the hydrophobic side chains of two consecutive residues of the peptide and the S1 and S2 pockets within the interdomain tunnel (Fig. 5a). When all PrfA-peptide structures were superimposed (including also monomer B of the PrfA-RGLL complex), it became apparent that the additional contacts with the S1 and S2 sites, are a general feature of the binding mechanism (Fig. 5b). At the S1 site, the sidechains of peptide residues Leu, Val and Pro bound. These amino acids made van der Waal’s interactions with the side chains of the aromatic residues Tyr63, Phe67, Tyr126, and Trp224. At the S2 site, the side chains of peptide residues Leu, Phe and the hydrophopic part of Arg made van der Waal’s interactions with the side chains of Ile45, Tyr62, Gln146, Ile149, Leu150, Tyr154, and Leu174 (Figs. 5b).

**Fig. 5.**
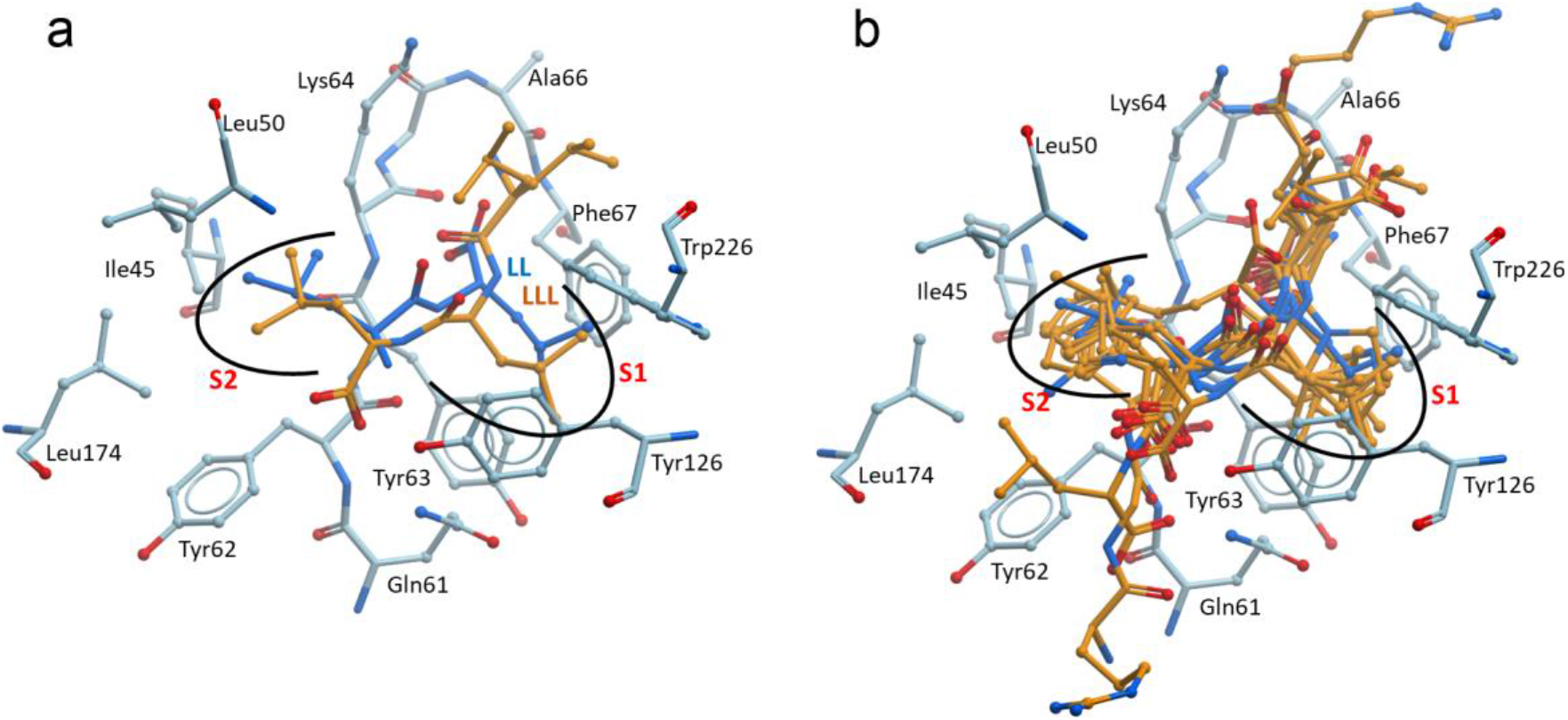
Peptide binding selectivity in PrfA. (**a**) Two subsequent leucine residues of the peptides LL and LLL bind at the S1 and S2 sites of the protein, which provide selectivity to the binding. (**b**) Superimposition of all PrfA-peptide complexes highlights that hydrophobic residues bound at the S1 and S2 sites are a common feature of peptide binding to PrfA.

The significance of these hydrophobic interactions at the S1 and S2 pockets is highlighted by how the TKPR peptide binds to PrfA (Supplementary Fig. 2). TKPR is comparatively more polar than the other peptides and only forms a single main-chain hydrogen bond to the protein. While only the positions of the Pro and Arg side chains could be modelled, their hydrophobic moieties are positioned at the S1 and S2 sites, respectively (Supplementary Fig. 2). Thus, parallel and antiparallel main-chain-main-chain contacts with PrfA β5 provide flexible sequence-independent peptide binding. Binding selectivity is however provided by two adjacent residues in the inhibitory peptides, capable of establishing hydrophobic contacts with the S1 and S2 pockets at the PrfA interdomain tunnel. The main chain atoms of these two hydrophobic residues make hydrogen bonds to the main chain atoms of Tyr62 and Lys64 positioned at PrfA β5. Their flanking residues are then positioned to make further contacts with PrfA, via hydrogen bonds to the main chain atoms of Leu60 and Gly65 or to the side chains of Gln61, Lys122, Tyr126, Lys130 and Gln146 (Supplementary Fig. 2).

### GSH and the inhibitory peptides differ in their binding to PrfA

The PrfA-activating co-factor GSH also binds at the interdomain tunnel site of PrfA. There it forms β-strand interactions with β5^29^ in a similar manner to the inhibitory peptides (^25^ and this study). The ITC and BLI data presented here also show that GSH and peptide inhibitors compete for the same site (Fig. 3 and Supplementary Fig. 2. However, even though the binding sites of inhibitory peptides and the GSH activator overlap, superimposing the structures reveals several fundamental differences that explain their different effects on PrfA activity. First, GSH is not a true peptide but a peptide-like molecule—a tripeptide with a gamma peptide linkage between the carboxyl group of the glutamate side chain and cysteine. As a consequence the distance between the N-terminal nitrogen and oxygen atoms of the gamma peptide linkage is 4.8 Å in GSH, whereas it is 2.7 Å in a regular residue, e.g., as seen in EVF (Fig. 6ab). This longer distance allows GSH to form more favourable hydrogen bond distances and angles to the main-chain oxygen atoms of Lys64 and Ala66, without disrupting the hydrogen bonding pattern to the rest of β5 (Fig. 6b). Second, GSH binding allows a structural movement of the C-terminal domain that closes the tunnel site and enables the formation of an active HTH motif. This movement positions critical amino acids (i.e. Gln146, Tyr154, and Leu174) in the vicinity of the HTH motif in the PrfA-GSH complex. In their new positions, these residues bridge the bound GSH molecule and the HTH motif, allowing a network of water molecules to form and connect the C-terminal carboxyl group of GSH to a now properly folded, and DNA-binding-compatible, HTH motif^29^. In other words, the unique set of interactions of GSH within the interdomain tunnel of PrfA leads to correct positioning of the DNA-binding helices into the perfect symmetry required for binding onto the dyad symmetrical target DNA sequence^12,29^. In particular, the position of the glycine residue of GSH at site S2 appears to be crucial for activation (Fig. 6b). This is because only glycine, lacking a side chain, can accommodate the bulky side chain of the incoming Tyr154 residue^29^. In contrast, bound inhibitor peptides with side chains positioned at S2, sterically prevent activating movements of Tyr154 and the rest of the C-terminal domain (Fig. 6c,d).

**Fig. 6.**
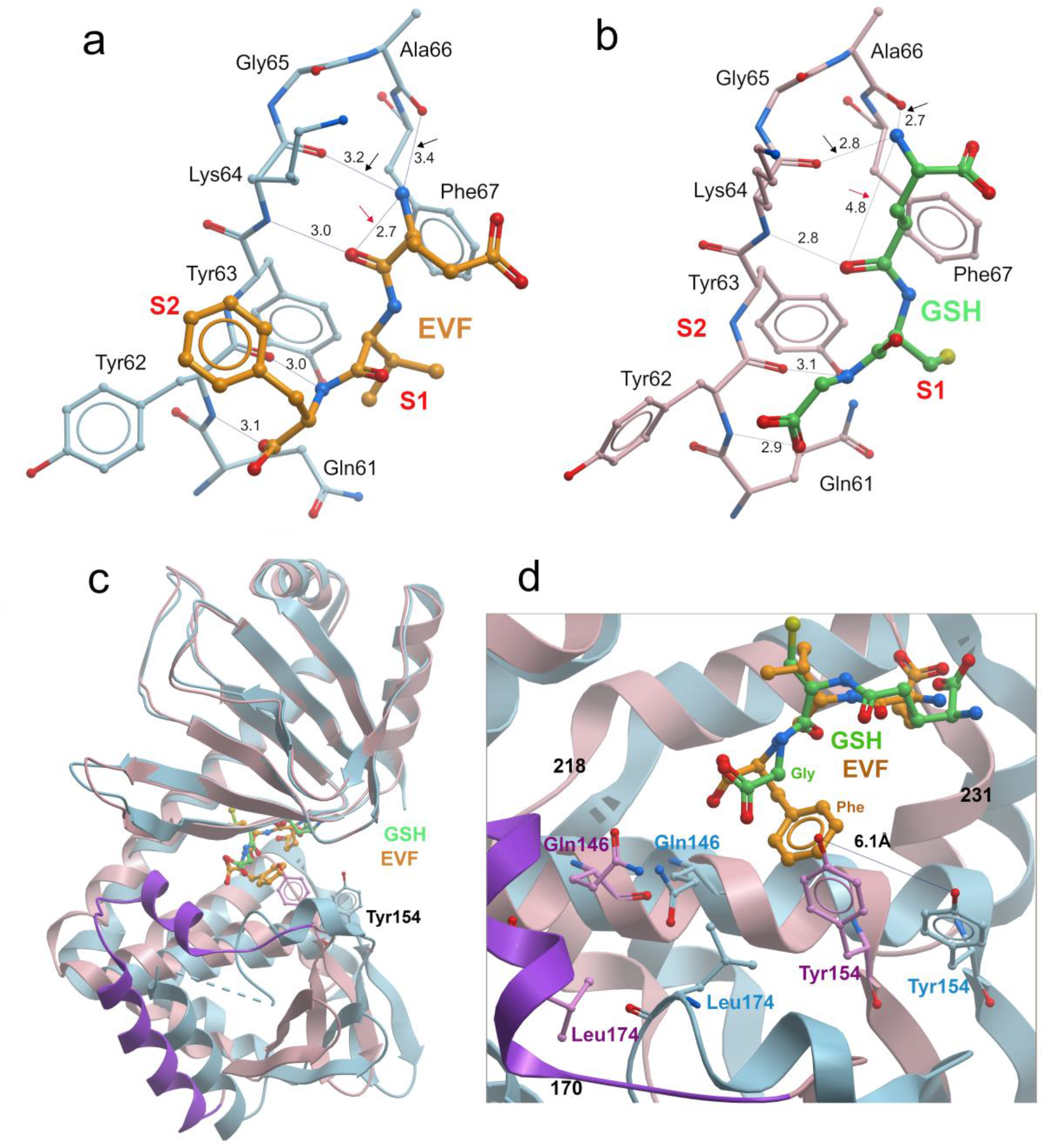
Differences in binding of the inhibitor peptide EVF (a) and the activator GSH molecule (b). The distance between the main chain nitrogen and oxygen atoms in the gamma peptide linkage in GSH is 2 Å longer than the same distance in a regular peptide (4.8 Å in GSH compared to 2.7 Å in EVF, red arrows). This enables GSH to make hydrogen bonds to residues Lys64 and Ala66 of β5 with favourable bonding distances and angles (black arrows). Furthermore, GSH has no hydrophobic side chain bound at S2, a common feature of all inhibitory peptides. For EVF, the side chain of Phe3 is positioned in site S2. Structural coordinates for the PrfA-GSH complex are from PDB code 5lrr^27^. (**c**) Superimposition of monomer A from EVF inhibited (blue) and GSH activated (pink) PrfA based on residues 2-123 in the N-terminal domain. Residue Tyr154 in both structures are shown as balls-and-sticks. (d) Close up view of the ligand binding site in (c) highlighting the closing in of the C-terminal domain of PrfA towards the GSH activator molecule at the tunnel site. Highlighted is also the steric clash between the Phe3 side chain of the EVF inhibitor and the Tyr154 side chain in the GSH activated structure (pink).

## Discussion

Protein-peptide interactions are ubiquitous in biological systems and mediate key cellular processes such as signalling and regulation, protein trafficking, DNA repair, or immune recognition^33-35^. Here we report the structural basis of a protein-peptide interaction that controls the activity of a transcriptional activator, PrfA, the master regulator of *Listeria* virulence^13^. The interacting peptides are small nutritional oligopeptides of different sequence and physicochemical properties scavenged from the medium via the listerial Opp transporter^25^. In this study, the mechanism of promiscuous binding of these peptides to PrfA was characterized.

Our structural analyses of PrfA-peptide interactions show that PrfA-peptide binding involves a combination of variability and selectivity. The variability is provided by unspecific main-chain–main-chain interactions (β-sheet-like), whereas the selectivity relies on hydrophobic interactions at two selectivity sites, S1 and S2^32^ found at the PrfA interdomain tunnel. In the tested peptides, these interactions involve residues Leu, Val, and Pro at the S1 site, and Leu, Phe, and the aliphatic side chain of Arg, at the S2 site. We suggest that these sites are important for defining the peptide’s binding affinity to PrfA. In addition, we anticipate that combinations of two sequential hydrophobic residues can bind at the S1 and S2 sites. Furthermore, the PrfA-TEPL structure showed binding of the tetrapeptide only to monomer A. This feature was also seen in the previously studied Prfa-LL dipeptide structure^25^ as well as in PrfA structures in complex with 2-pyridones inhibitors ^31^. This suggests the presence of allosteric communications within the PrfA dimer, in which binding of molecules in the tunnel site of one monomer inhibits binding of a second compound in the partner monomer.

PrfA-peptide binding bears some of the general features of protein-peptide interactions, reinforcing the notion that common principles underlie the binding of intrinsically flexible peptide ligands to proteins^35^. These include: preferential binding in β-strand/β-strand-like conformation, binding involving hydrogen bonds with the peptide backbone, and binding also generally relies on “hot-spot” or “anchor” residues in the peptide, with a predominance of leucine and aromatic or hydrophobic residues^33,35-38^, as seen in our case.

Peptide-mediated PrfA inhibition by competitive occupancy of the tunnel binding site for GSH^25,29^, itself a tripeptide, could be attributed to rapid, transient protein-peptide interactions with relatively weak affinity, like those that dynamically regulate cellular processes via Short Linear Motifs (SLiMs)^39^. Indeed, the *K*_d_ values for PrfA-inhibitory peptides (previously estimated to ≈25 μM^25^ and in this study between 2-145 μM as determined with ITC, exceptions being TKPR and TEPL whose *K*_d_ values could not be determined with our experimental setup) are in the range of those determined for SLiMs and their protein targets (*K*_d_ of 1-500 μM). This is opposed to the nM range of highly specific ligand-receptor interactions^39^. Whereas peptide and SLiMs-protein interactions are mostly specific^35,39,40^, a salient feature of peptide-mediated PrfA inhibition is its promiscuity^25^. The two major determinants of this binding promiscuity, as revealed by our study, are probably the unique ability of the inhibitory peptides to interact with PrfA β5 in both parallel and antiparallel conformations, and the non-specific accommodation of hydrophobic and aromatic residues of the peptide in the spacious pockets within the PrfA interdomain tunnel.

Additionally, oligopeptides of different sizes, including octapeptides, can inhibit PrfA-dependent gene expression^25^. While longer peptides may exert their effects when metabolically processed into smaller peptides^25^, it is evident from our PrfA-peptide structures that peptides longer than four residues can also directly inhibit PrfA. As shown in Fig. 7, we anticipate that the flanking residues of peptides equal to or longer than six-mers are fully exposed on the surface of the PrfA protein, with the centrally positioned hydrophobic residues buried in the tunnel and the terminal residues exposed at both interdomain tunnel entrances. Whether the overhanging residues can modulate binding to PrfA by interacting with specific features at either entrance of the interdomain tunnel remains to be determined.

**Fig. 7.**
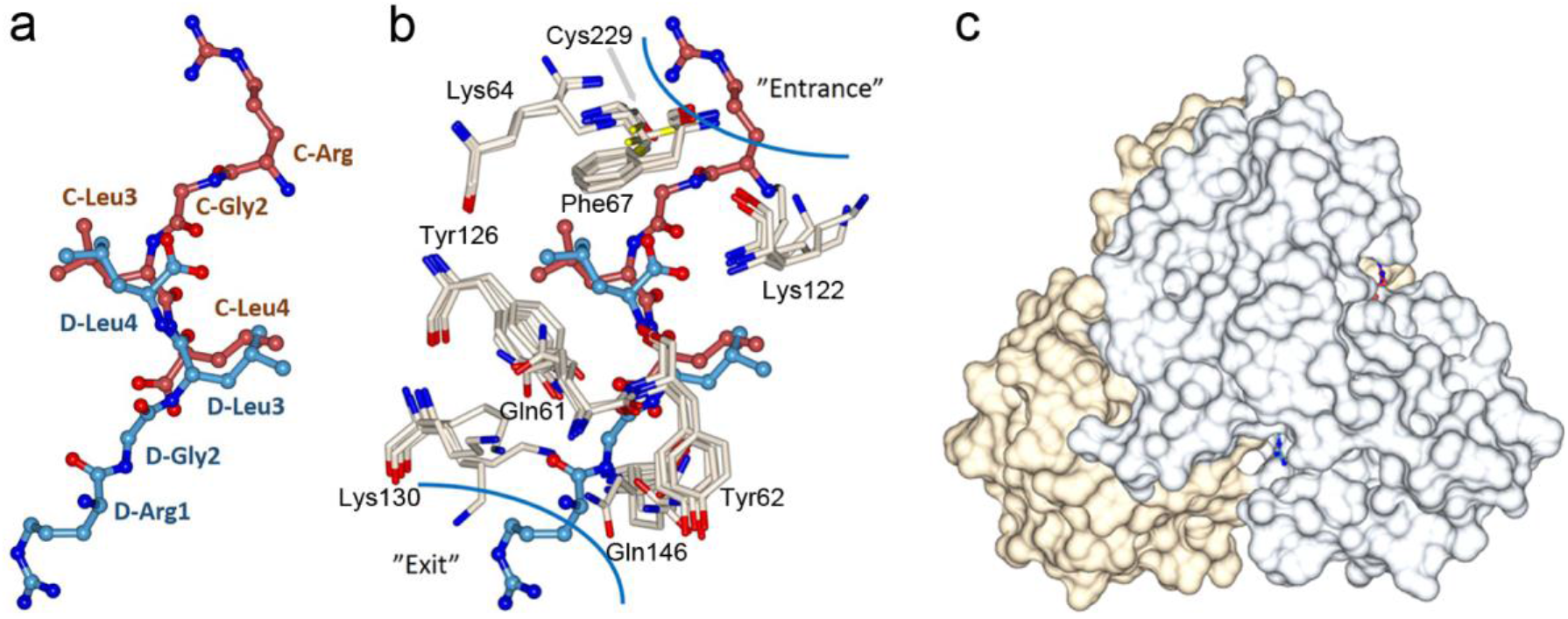
Gatekeeper residues at the tunnel site. (**a**) Superimposition of monomers A and B in the PrfA-RGLL complex positions parallel and antiparallel RGLL peptides into one artificial RGLLGR peptide. The figure shows C-RGLL (red) and D-RGLL (blue) bound in monomers A and B, respectively. Combined the peptides stretch over 6 residues. (**b**) Gatekeeper residues, i.e. Tyr62, Lys122, Lys 130 and Gln146 can change their rotamers to accommodate variable peptide side chains at the entrance and exit of the tunnel. For clarity, only residues from PrfA bound to LLL in monomer A, EVF in monomer A, and RGLL in monomers A and B are shown. (**c**) The length and position of the artificial RGLLGR peptide suggests that, for example, hexapeptides will have their first and last residues protruding out of the tunnel site if residues 3 and 4 are hydrophobic and bind at the S1 and S2 sites.

The broad binding specificity of peptides to PrfA mirrors the sequence-independent peptide binding of the bacterial ABC oligopeptide transporters, one of which is the listerial Opp permease involved in the uptake of PrfA-activating and -inhibiting peptides^25^. The peptide receptor subunit of these transporters—OppA and homologs such as AppA from *Bacillus* or DppA in gram-negative bacteria^41-45^—shares striking similarities with the binding mechanism in PrfA. This is likely a consequence of convergent evolution towards broad specificity in peptide recognition. Thus, the bound peptide substrates in OppA are in β-strand conformation, although in OppA they make main-chain contacts with the protein on both sides (Supplementary Fig. 4a and b). The bound peptides are similarly buried in a cavity between two large receptor protein lobes, where water-filled hydrophobic pockets can readily accommodate diverse side chains imposing little binding specificity^25,27^. At the same time, they preferentially bind to hydrophobic residues defining the peptide’s binding register. The peptides also bind with similar affinity to the PrfA-inhibitory peptides. In addition, PrfA and the peptide receptor subunits of the oligopeptide transporters share the ability to bind to small di- and tri-peptides^25,44-47^.

The regulation of PrfA activity by small exogenous oligopeptides from the bacterial environment is reminiscent Gram-positive quorum sensing mechanisms, involving Opp-transported (re-imported) peptide pheromones and short hydrophobic peptides (SHP)^48^. In these systems, small 5- to 8-residue signalling peptides directly bind to transcription factors, the so-called RRNPP family of peptide-sensing regulators^49,50^. These interactions typically occur in deep interdomain clefts and involve hydrogen bonding of the extended peptide backbone to the regulator, and non-bonded hydrophobic interactions, as well as hydrogen bonds, with the peptide side chains confer specificity to the interaction^51-54^. However, unlike PrfA, these interactions often lead to substantial conformational changes from inactive monomeric to active multimeric forms^51-54^ and have binding affinities that are two orders of magnitude higher (K_d_ ≈0.1-0.5 μM)^51,55,56^ compared to those of PrfA and its inhibitory peptides.

The RRNPP regulators and now the listerial virulence regulator PrfA are paradigmatic examples of how peptides mediate the control of transcription factor activity. RRNPP quorum-sensing signalling peptides can exhibit permissive recognition properties within a given regulator subfamily, enabling bacterial interspecies communication^55,57,58^. PrfA is however unique in its very broad peptide binding specificity. This mechanism allows the facultative pathogen *L. monocytogenes* to respond to peptide signals of diverse origins—animal, plant, microbial, and even self-derived to fine-tune the costly expression of its virulence factors in response to environmental cues^5,25^.

PrfA activity is essential for the pathogenicity of *L. monocytogenes* and is therefore an obvious target for the development of anti-infectives. Previous studies identified a group of ring-fused 2-pyridone heterocycles that inhibit PrfA and reduce virulence factor expression in *L. monocytogenes*^31,32^. Two groups on the pyridone scaffold were found to be important for inhibition: a small hydrophobic group that binds to the PrfA tunnel site S1, and a second, larger hydrophobic group, based on a naphtyl group, for binding to site S2. Thus, peptides and ring-fused 2-pyridone inhibitors bind to the PrfA interdomain tunnel in a similar manner despite their structural differences. The structural insights from the PrfA-peptide complexes presented here may guide further development of anti-virulence precision anti-infectives targeting PrfA.

## Methods

### Characterization of PrfA peptide inhibitors

PrfA inhibition studies were performed using *L. monocytogenes* P14-P_hly-lux_ as previously described^25^. P14-P_hly-lux_ is a wild-type serovar 4b isolate carrying a chromosomally integrated *luxABCDE* reporter under the control of the PrfA-regulated *hly* promoter. Peptides were added at 1 mM concentration to PrfA-activating chemically defined medium (CDM)^25^.

#### Protein purification

For crystallization studies, PrfA was recombinantly expressed in *E. coli* using the pET28a with a 6-His tag and tobacco etch virus (TEV) cleavage site. The construct encoded the full-length PrfA^WT^ protein with two non-native N-terminal residues (GA) at TEV cleavage site. The cleaved product was purified on a MonoS 5/5 ion-exchange column (GE-Healthcare) with elution at ∼250 mM NaCl in 10 mM Tris pH 7.5, 1 mM DTT. This was followed by size-exclusion chromatography on a HiLoad Superdex 75 16/60 column (GE Healthcare) equilibrated in 20 mM sodium phosphate pH 6.5, 200 mM NaCl. The fractions containing PrfA were pooled and concentrated using a Centriprep-10 centrifugal concentrator (Millipore).

#### Isothermal titration calorimetry

PrfA and oligopeptides were prepared in a buffer containing 20 mM Tris-HCl pH 8 and 200 mM NaCl. In the calorimetry experiments, 50 μM PrfA was titrated against 0.5 mM or 2 mM oligopeptides. The experiments were performed at 25°C using a MicroCal auto-ITC200 calorimeter (MicroCal-Malvern) and the standard methods “Plates Prerinse Syringe Clean” And “EDTA”. After subtracting the isotherms obtained by injecting oligopeptides into the buffer alone, the binding isotherms were fitted to the “one set of sites” model implemented in Origin 7 (OriginLab). The experiments were performed in independent triplicate experiments.

#### Bio-layer interferometry

The binding and dissociation of PrfA to DNA was measured in real time using an Octet system (Sartorius). For this purpose, a 5’-biotinylated DNA oligo (Eurofins) containing the P*plcA*/P*hly* PrfA box was annealed with the non-biotinylated complementary DNA oligo by mixing in a 1:1.1 ratio, heating the mixture for 5 min in boiling water and slow cooling.

P*plcA*/P*hly*-F: 5’-[BIOTEG]TGTCCCTTTATCGTCGTTAACAAATGTTAATGCCTCGACA-3’ P*plcA*/P*hly*-R: 5’-TGTCGAGGCATTAACATTTGTTAACGACGATAAAGGGACA-3’

100 nM dsDNA was captured on Streptavidin (SA) biosensors and incubated at 25°C with a shaking speed of 1000 rpm in a buffer containing 50 mM Tris-HCl pH 7.5, 300 mM NaCl, and 0.05% Tween-20. For oligopeptide-inhibition experiments, biosensors were incubated in a twofold dilution series of PrfA-oligopeptide mixtures (1:100 molar ratio) and a PrfA-GSH mixture (GSH maintained at 5 mM) in the same buffer. For GSH-competition experiments, 5 mM GSH was added to a mixture of 5 µM PrfA and 500 µM oligopeptides in the same buffer. All experiments were performed independently in triplicate. The times for the baseline, association, and dissociation steps were 180, 300, and 600 s, respectively. After subtracting the control and reference sensor signals, the sensorgrams were analysed using Octet Analysis Studio software to determine the binding responses.

#### Crystallization and data collection

For the co-crystallization, 3.1-6.6 mg mL^−1^ of PrfA in 20 mM sodium phosphate at pH 6.5, 200 mM NaCl was mixed with the peptides and dithiothreitol (DTT) to give a 1:10 molar ratio of protein to peptide and final concentrations of 0.5-2.5 mM peptides, 0.5-2.5% (v/v) DMSO and 1-4 mM DTT. The mixture was incubated at room temperature or 4°C for 4 h to 14 h before crystallization setup. Crystallization was performed by the hanging drop or sitting drop vapor diffusion method at 18°C. For this, the protein-peptide mixture was mixed in a 1:1 ratio with the crystallization solution containing 100 mM Na citrate at pH 5.1 to 5.7, 16 % (w/v) to 26 % (w/v) PEG4000 and 0 % (v/v) or 17 % (v/v) isopropanol. All crystals, except for the PrfA-tri-leucine complex (PrfA-LLL), were additionally soaked with 20 mM peptides, 35% (w/v) PEG4000 and 100 mM sodium citrate before flash-freezing in liquid nitrogen. The cryoprotection solution consisted of the corresponding crystallization condition and an increased PEG4000 concentration of 35 % (w/v). Exact crystallization conditions are given in the Protein Data Bank for each PrfA-peptide complex, Data collection was performed at 100 K at synchrotrons specified in Supplementary Table 1.

#### Processing, phasing and refinement

Diffraction images were processed with XDS^59^ and scaled and merged using AIMLESS from the CCP4 software suite^60,61^. The structure was determined by molecular replacement with the PHASER program from the PHENIX program suite^62,63^ using the wild-type PrfA-di-leucine complex structure (PrfA-LL, PDB code 6hck^25^) as the search model. The atomic models were manually built^64^ and refined with PHENIX Refine^65^. Noncrystallographic symmetry restraints were not applied during refinement. The quality of the electron density map of the ligand was significantly improved in the POLDER omit map^30^, and the ligand was modelled with the aid of LigandFit^66,67^. The recognition helices were flexible and could not be modelled (∼ residues 174-182) for all structures. In addition, the C-terminal DNA binding domain was poorly defined in the electron density of monomer A of the PrfA-TEPL complex structures, meaning residues 198-208 could not be modelled. Furthermore, the side chain of residue Glu2 in the TEPL peptide was not defined in the electron density. Refinement statistics are shown in Supplementary Table 1. Figures were prepared with CCP4mg^68^ or ICM browser (Molsoft LLC).

## Data availability

The atomic coordinates and the structure factors have been deposited with the Protein Data Bank^69^ (PDB codes 8cb4 for PrfA-LLL, 8cb5 for PrfA-EVF, 8cb7 for PrfA-EVFL, 8cb8 for PrfA-STLL, 8cbg for PrfA-RGLL, 8cbi for PrfA-TKPR, and 8cbp for PrfA-TEPL).

## Author contributions

T.H., C.L., and E.S.-E. performed X-ray crystallographic studies of the PrfA-peptide complexes. C.G. purified the PrfA protein. M.S and E.K performed peptide inhibition studies. J.V-B and E.S.-E. conceived the study, designed research, acquired funding, and wrote the paper. All authors had editorial input on the final version of the manuscript

## Acknowledgments

ESA was funded by the Swedish Research Council (ID 2015-03607 and 2019-03771). In addition, funds are gratefully acknowledged from the Kempe Foundation and the Erling-Persson Family Foundation. Work at the JV-B laboratory was supported by Wellcome (program grant WT074020MA) and core Roslin Institute Strategic Programme funding from the BBSRC (BB/J004227/1). We thank beamline staff at the European Synchrotron Radiation Facility (ESRF, France) and the Swiss Light Source (SLS, Switzerland) for support and access to beamlines ID23-2 and X06DA, respectively. We acknowledge MAX IV Laboratory and staff for time on Beamline Biomax under proposal 20180236. Research conducted at MAX IV, a Swedish national user facility, is supported by the Swedish Research council under contract 2018-07152, the Swedish Governmental Agency for Innovation Systems under contract 2018-04969, and Formas under contract 2019-02496.

## References

1. Freitag, N.E., Port, G.C. & Miner, M.D. Listeria monocytogenes - from saprophyte to intracellular pathogen. Nat Rev Microbiol 7, 623–8 (2009).

2. Radoshevich, L. & Cossart, P. Listeria monocytogenes: towards a complete picture of its physiology and pathogenesis. Nat Rev Microbiol 16, 32–46 (2018).

3. Koopmans, M.M., Brouwer, M.C., Vazquez-Boland, J.A. & van de Beek, D. Human Listeriosis. Clin Microbiol Rev, e0006019 (2022).

4. Sturm, A. et al. The cost of virulence: retarded growth of Salmonella Typhimurium cells expressing type III secretion system 1. PLoS Pathog 7, e1002143 (2011).

5. Vasanthakrishnan, R.B. et al. PrfA regulation offsets the cost of Listeria virulence outside the host. Environ Microbiol 17, 4566–79 (2015).

6. Tiensuu, T., Guerreiro, D.N., Oliveira, A.H., O’Byrne, C. & Johansson, J. Flick of a switch: regulatory mechanisms allowing Listeria monocytogenes to transition from a saprophyte to a killer. Microbiology (Reading) 165, 819–833 (2019).

7. Chakraborty, T. et al. Coordinate regulation of virulence genes in Listeria monocytogenes requires the product of the prfA gene. J Bacteriol 174, 568–74 (1992).

8. Freitag, N.E., Rong, L. & Portnoy, D.A. Regulation of the prfA transcriptional activator of Listeria monocytogenes: multiple promoter elements contribute to intracellular growth and cell-to-cell spread. Infect Immun 61, 2537–44 (1993).

9. Mengaud, J. et al. Pleiotropic control of Listeria monocytogenes virulence factors by a gene that is autoregulated. Mol Microbiol 5, 2273–83 (1991).

10. Leimeister-Wachter, M., Haffner, C., Domann, E., Goebel, W. & Chakraborty, T. Identification of a gene that positively regulates expression of listeriolysin, the major virulence factor of listeria monocytogenes. Proc Natl Acad Sci U S A 87, 8336–40 (1990).

11. Hamon, M., Bierne, H. & Cossart, P. Listeria monocytogenes: a multifaceted model. Nat Rev Microbiol 4, 423–34 (2006).

12. Scortti, M., Monzo, H.J., Lacharme-Lora, L., Lewis, D.A. & Vazquez-Boland, J.A. The PrfA virulence regulon. Microbes Infect 9, 1196–207 (2007).

13. de las Heras, A., Cain, R.J., Bielecka, M.K. & Vazquez-Boland, J.A. Regulation of Listeria virulence: PrfA master and commander. Curr Opin Microbiol 14, 118–27 (2011).

14. Johansson, J. & Freitag, N.E. Regulation of Listeria monocytogenes Virulence. Microbiol Spectr 7(2019).

15. Toledo-Arana, A. et al. The Listeria transcriptional landscape from saprophytism to virulence. Nature 459, 950–6 (2009).

16. Johansson, J. et al. An RNA thermosensor controls expression of virulence genes in Listeria monocytogenes. Cell 110, 551–61 (2002).

17. Park, S.F. & Kroll, R.G. Expression of listeriolysin and phosphatidylinositol-specific phospholipase C is repressed by the plant-derived molecule cellobiose in Listeria monocytogenes. Mol Microbiol 8, 653–61 (1993).

18. Brehm, K., Ripio, M.T., Kreft, J. & Vazquez-Boland, J.A. The bvr locus of Listeria monocytogenes mediates virulence gene repression by beta-glucosides. J Bacteriol 181, 5024–32 (1999).

19. Dos Santos, P.T., Thomasen, R.S.S., Green, M.S., Faergeman, N.J. & Kallipolitis, B.H. Free Fatty Acids Interfere with the DNA Binding Activity of the Virulence Regulator PrfA of Listeria monocytogenes. J Bacteriol 202(2020).

20. Haber, A. et al. L-glutamine Induces Expression of Listeria monocytogenes Virulence Genes. PLoS Pathog 13, e1006161 (2017).

21. Lobel, L. et al. The metabolic regulator CodY links Listeria monocytogenes metabolism to virulence by directly activating the virulence regulatory gene prfA. Mol Microbiol 95, 624–44 (2015).

22. Gaballa, A., Guariglia-Oropeza, V., Wiedmann, M. & Boor, K.J. Cross Talk between SigB and PrfA in Listeria monocytogenes Facilitates Transitions between Extra- and Intracellular Environments. Microbiol Mol Biol Rev 83(2019).

23. Reniere, M.L. et al. Glutathione activates virulence gene expression of an intracellular pathogen. Nature 517, 170–3 (2015).

24. Gopal, S. et al. A multidomain fusion protein in Listeria monocytogenes catalyzes the two primary activities for glutathione biosynthesis. J Bacteriol 187, 3839–47 (2005).

25. Krypotou, E. et al. Control of Bacterial Virulence through the Peptide Signature of the Habitat. Cell Rep 26, 1815–1827 e5 (2019).

26. Vega, Y. et al. New Listeria monocytogenes prfA* mutants, transcriptional properties of PrfA* proteins and structure-function of the virulence regulator PrfA. Mol Microbiol 52, 1553–65 (2004).

27. Eiting, M., Hageluken, G., Schubert, W.D. & Heinz, D.W. The mutation G145S in PrfA, a key virulence regulator of Listeria monocytogenes, increases DNA-binding affinity by stabilizing the HTH motif. Mol Microbiol 56, 433–46 (2005).

28. Sheehan, B., Klarsfeld, A., Msadek, T. & Cossart, P. Differential activation of virulence gene expression by PrfA, the Listeria monocytogenes virulence regulator. J Bacteriol 177, 6469–76 (1995).

29. Hall, M. et al. Structural basis for glutathione-mediated activation of the virulence regulatory protein PrfA in Listeria. Proc Natl Acad Sci U S A 113, 14733–14738 (2016).

30. Liebschner, D. et al. Polder maps: improving OMIT maps by excluding bulk solvent. Acta Crystallogr D Struct Biol 73, 148–157 (2017).

31. Good, J.A. et al. Attenuating Listeria monocytogenes Virulence by Targeting the Regulatory Protein PrfA. Cell Chem Biol 23, 404–14 (2016).

32. Kulen, M. et al. Structure-Based Design of Inhibitors Targeting PrfA, the Master Virulence Regulator of Listeria monocytogenes. J Med Chem 61, 4165–4175 (2018).

33. Stanfield, R.L. & Wilson, I.A. Protein-peptide interactions. Curr Opin Struct Biol 5, 103–13 (1995).

34. Pawson, T. & Nash, P. Assembly of cell regulatory systems through protein interaction domains. Science 300, 445–52 (2003).

35. London, N., Movshovitz-Attias, D. & Schueler-Furman, O. The structural basis of peptide-protein binding strategies. Structure 18, 188–99 (2010).

36. Eisen, H.N. et al. Promiscuous binding of extracellular peptides to cell surface class I MHC protein. Proc Natl Acad Sci U S A 109, 4580–5 (2012).

37. Bhattacherjee, A. & Wallin, S. Exploring Protein-Peptide Binding Specificity through Computational Peptide Screening. PLoS Comput Biol 9, e1003277 (2013).

38. Vanhee, P. et al. Protein-peptide interactions adopt the same structural motifs as monomeric protein folds. Structure 17, 1128–36 (2009).

39. Nguyen, H.Q. et al. Quantitative mapping of protein-peptide affinity landscapes using spectrally encoded beads. Elife 8(2019).

40. Parker, B.W. et al. Mapping low-affinity/high-specificity peptide-protein interactions using ligand-footprinting mass spectrometry. Proc Natl Acad Sci U S A 116, 21001–21011 (2019).

41. Monnet, V. Bacterial oligopeptide-binding proteins. Cell Mol Life Sci 60, 2100–14 (2003).

42. Tame, J.R. et al. The structural basis of sequence-independent peptide binding by OppA protein. Science 264, 1578–81 (1994).

43. Dunten, P. & Mowbray, S.L. Crystal structure of the dipeptide binding protein from Escherichia coli involved in active transport and chemotaxis. Protein Sci 4, 2327–34 (1995).

44. Berntsson, R.P. et al. The structural basis for peptide selection by the transport receptor OppA. EMBO J 28, 1332–40 (2009).

45. Levdikov, V.M. et al. The structure of the oligopeptide-binding protein, AppA, from Bacillus subtilis in complex with a nonapeptide. J Mol Biol 345, 879–92 (2005).

46. Tame, J.R., Dodson, E.J., Murshudov, G., Higgins, C.F. & Wilkinson, A.J. The crystal structures of the oligopeptide-binding protein OppA complexed with tripeptide and tetrapeptide ligands. Structure 3, 1395–406 (1995).

47. Berntsson, R.P., Thunnissen, A.M., Poolman, B. & Slotboom, D.J. Importance of a hydrophobic pocket for peptide binding in lactococcal OppA. J Bacteriol 193, 4254–6 (2011).

48. Slamti, L. & Lereclus, D. The oligopeptide ABC-importers are essential communication channels in Gram-positive bacteria. Res Microbiol 170, 338–344 (2019).

49. Rocha-Estrada, J., Aceves-Diez, A.E., Guarneros, G. & de la Torre, M. The RNPP family of quorum-sensing proteins in Gram-positive bacteria. Appl Microbiol Biotechnol 87, 913–23 (2010).

50. Declerck, N. et al. Structure of PlcR: Insights into virulence regulation and evolution of quorum sensing in Gram-positive bacteria. Proc Natl Acad Sci U S A 104, 18490–5 (2007).

51. Zouhir, S. et al. Peptide-binding dependent conformational changes regulate the transcriptional activity of the quorum-sensor NprR. Nucleic Acids Res 41, 7920–33 (2013).

52. Talagas, A. et al. Structural Insights into Streptococcal Competence Regulation by the Cell-to-Cell Communication System ComRS. PLoS Pathog 12, e1005980 (2016).

53. Neiditch, M.B., Capodagli, G.C., Prehna, G. & Federle, M.J. Genetic and Structural Analyses of RRNPP Intercellular Peptide Signaling of Gram-Positive Bacteria. Annu Rev Genet 51, 311–333 (2017).

54. Do, H. & Kumaraswami, M. Structural Mechanisms of Peptide Recognition and Allosteric Modulation of Gene Regulation by the RRNPP Family of Quorum-Sensing Regulators. J Mol Biol 428, 2793–804 (2016).

55. Shanker, E. et al. Pheromone Recognition and Selectivity by ComR Proteins among Streptococcus Species. PLoS Pathog 12, e1005979 (2016).

56. Aggarwal, C., Jimenez, J.C., Nanavati, D. & Federle, M.J. Multiple length peptide-pheromone variants produced by Streptococcus pyogenes directly bind Rgg proteins to confer transcriptional regulation. J Biol Chem 289, 22427–36 (2014).

57. Fleuchot, B. et al. Rgg-associated SHP signaling peptides mediate cross-talk in Streptococci. PLoS One 8, e66042 (2013).

58. Cook, L.C. & Federle, M.J. Peptide pheromone signaling in Streptococcus and Enterococcus. FEMS Microbiol Rev 38, 473–92 (2014).

59. Kabsch, W. Xds. Acta Crystallogr D Biol Crystallogr 66, 125–32 (2010).

60. Evans, P.R. & Murshudov, G.N. How good are my data and what is the resolution? Acta Crystallogr D Biol Crystallogr 69, 1204–14 (2013).

61. Winn, M.D. et al. Overview of the CCP4 suite and current developments. Acta Crystallogr D Biol Crystallogr 67, 235–42 (2011).

62. McCoy, A.J. et al. Phaser crystallographic software. J Appl Crystallogr 40, 658–674 (2007).

63. Liebschner, D. et al. Macromolecular structure determination using X-rays, neutrons and electrons: recent developments in Phenix. Acta Crystallogr D Struct Biol 75, 861–877 (2019).

64. Emsley, P., Lohkamp, B., Scott, W.G. & Cowtan, K. Features and development of Coot. Acta Crystallogr D Biol Crystallogr 66, 486–501 (2010).

65. Afonine, P.V. et al. Towards automated crystallographic structure refinement with phenix.refine. Acta Crystallogr D Biol Crystallogr 68, 352–67 (2012).

66. Terwilliger, T.C., Klei, H., Adams, P.D., Moriarty, N.W. & Cohn, J.D. Automated ligand fitting by core-fragment fitting and extension into density. Acta Crystallogr D Biol Crystallogr 62, 915–22 (2006).

67. Terwilliger, T.C., Adams, P.D., Moriarty, N.W. & Cohn, J.D. Ligand identification using electron-density map correlations. Acta Crystallogr D Biol Crystallogr 63, 101–7 (2007).

68. McNicholas, S., Potterton, E., Wilson, K.S. & Noble, M.E. Presenting your structures: the CCP4mg molecular-graphics software. Acta Crystallogr D Biol Crystallogr 67, 386–94 (2011).

69. Berman, H.M. et al. The Protein Data Bank and the challenge of structural genomics. Nat Struct Biol 7 Suppl, 957–9 (2000).

