## Supplementary Data for "Structural basis of promiscuous inhibition of *Listeria* virulence activator PrfA by oligopeptides"

### Table S1. Data collection and refinement statistics

| | PrfA-LLL | PrfA-EVF | PrfA-EVFL |
|---|---|---|---|
| **Data collection** | | | |
| Synchrotron | ESRF, Grenoble, France | MaxIV, Lund, Sweden | SLS, Switzerland |
| Beam line | ID23-2 | Biomax | X06DA2 |
| Wavelength (Å) | 0.8731 | 0.9537 | 1.0000 |
| Space group | $P2_12_12_1$ | $P2_12_12_1$ | $P2_12_12_1$ |
| Cell dimensions[illegible] | | | |
| *a*, *b*, *c* (Å) | 48.10, 86.79, 115.86 | 48.20, 87.31, 112.64 | 48.62, 94.24, 101.55 |
| [illegible] (°) | 90.0, 90.0, 90.0 | 90.0, 90.0, 90.0 | 90.0, 90.0, 90.0 |
| Resolution (Å) | 48.1–2.57 (2.66–2.57)* | 47.3–2.25 (2.33–2.25)* | 44.7-2.85 (2.95-2.85) |
| $R_{merge}$ | 0.130 (1.182) | 0.260 (2.010) | 0.210 (2.001) |
| $R_{PIM}$ | 0.099 (0.927) | 0.053 (0.590) | 0.107 (1.013) |
| *I* / [illegible]*I* | 8.0 (1.0) | 12.9 (1.3) | 7.7 (1.2) |
| Half-set correlation CC(1/2) | 0.99 (0.30) | 0.999 (0.546) | 0.993 (0.403) |
| Completeness (%) | 99.4 (98.1) | 100.0 (100.0) | 100.0 (100.0) |
| Redundancy | 4.6 (4.2) | 12.7 (13.4) | 8.6 (9.1) |
| Wilson B-factor (Å$^2$) | 53.1 | 53.6 | 69.9 |
| **Refinement** | | | |
| Resolution (Å) | 43.4-2.57 | 44.4-2.25 | 47.7-2.85 |
| No. reflections | 15932 (1521) | 23242 (2263) | 11417 (1113) |
| $R_{work}$ | 0.215 (0.300) | 0.225 (0.266) | 0.265 (0.36.2) |
| $R_{free}$ | 0.276 (0.353) | 0.305 (0.346) | 0.309 (0.397) |
| No. atoms | | | |
| Protein | 3762 | 3660 | 3720 |
| Peptides | 58 (LLL) | 56 (EVF) | 72 (EVFL) |
| Ligands¤ | 2 | 14 | 0 |
| Water | 36 | 53 | 3 |
| Clashscore | 2.9 | 3.5 | 7.4 |
| *B*-factors (Å$^2$) | | | |
| Protein | 66.1 | 69.5 | 77.9 |
| Peptides | 78.4 | 79.8 | 76.3 |
| Ligand/ion | 52.0 | 61.2 | - |
| Water | 50.0 | 56.7 | 43.6 |
| R.m.s. deviations | | | |
| Bond lengths (Å) | 0.005 | 0.009 | 0.010 |
| Bond angles (°) | 0.55 | 0.91 | 1.32 |
| PDB code | 8CB4 | 8CB5 | 8CB7 |

*One crystal was used for data collection. Values in parentheses are for the highest-resolution shell.
¤Ligands include: isopropanol, $PO_4^{3-}$, and $Na^+$. Estimate of resolution limit is based on CC1/2.

### Table S1. Data collection and refinement statistics (continued)

| | PrfA-STLL | PrfA-RGLL |
|---|---|---|
| **Data collection** | | |
| Synchrotron | MaxIV, Lund, Sweden | SLS, Switzerland |
| Beam line | Biomax | X06DA |
| Wavelength (Å) | 0.9184 | 1.0000 |
| Space group | $P2_12_12_1$ | $P2_1$ |
| Cell dimensions [illegible] | | |
| *a*, *b*, *c* (Å) | 47.95, 86.13, 115.22 | 55.52, 81.00, 60.46 |
| [illegible] (°) | 90.0, 90.0, 90.0 | 90.0, 113.18, 90.0 |
| Resolution (Å) | 48.0–2.25 (2.33–2.25)* | 48.2-3.00 (3.18-3.00) |
| $R_{merge}$ | 0.26 (4.411) | 0.15 (1.308) |
| $R_{PIM}$ | 0.106 (1.797) | 0.086 (0.808) |
| *I* / [illegible]*I* | 9.2 (1.6) | 6.7 (1.1) |
| Half-set correlation CC(1/2) | 0.996 (0.671) | 0.992 (0.431) |
| Completeness (%) | 100.0 (100.0) | 100.0 (100.0) |
| Redundancy | 13.4 (13.5) | 4.6 (4.4) |
| Wilson B-factor (Å$^2$) | 44.9 | 75.2 |
| **Refinement** | | |
| Resolution (Å) | 47.9-2.25 | 48.2-3.00 |
| No. reflections | 23371 (2282) | 9945 (978) |
| $R_{work}$ | 0.232 (0.289) | 0.231 (0.389) |
| $R_{free}$ | 0.286 (0.436) | 0.281 (0.462) |
| No. atoms | | |
| Protein | 3729 | 3772 |
| Peptides | 48 (-TLL) | 64 (RGLL) |
| Ligands¤ | 12 | 0 |
| Water | 57 | 7 |
| Clashscore | 4.0 | 4.8 |
| *B*-factors (Å$^2$) | | |
| Protein | 62.7 | 97.7 |
| Peptides | 73.8 | 103.6 |
| Ligand/ion | 96.1 | 0 |
| Water | 46.7 | 56.6 |
| R.m.s. deviations | | |
| Bond lengths (Å) | 0.004 | 0.003 |
| Bond angles (°) | 0.57 | 0.56 |
| PDB code | 8CB8 | 8CBG |

### Table S1. Data collection and refinement statistics (continued)

| | PrfA-TKPR | PrfA-TEPL |
|---|---|---|
| **Data collection** | | |
| Synchrotron | SLS, Switzerland | MaxIV, Lund, Sweden |
| Beam line | X06DA | Biomax |
| Wavelength (Å) | 1.0000 | 0.9537 |
| Space group | $P2_1$ | $P2_1$ |
| Cell dimensions [illegible] | | |
| *a*, *b*, *c* (Å) | 56.40, 80.95, 61.79 | 56.19, 81.39, 60.63 |
| [illegible] (°) | 90.0, 110.74, 90.0 | 90.0, 112.28, 90.0 |
| Resolution (Å) | 48.4–2.40 (2.49–2.40)* | 48.4-2.70 (2.83-2.70) |
| $R_{merge}$ | 0.149 (1.489) | 0.051 (0.876) |
| $R_{PIM}$ | 0.122 (1.225) | 0.047 (0.782) |
| *I* / [illegible]*I* | 5.6 (1.2) | 10.5 (1.2) |
| Half-set correlation CC(1/2) | 0.989 (0.300) | 0.997 (0.767) |
| Completeness (%) | 100.0 (100.0) | 99.7 (97.6) |
| Redundancy | 4.6 (4.7) | 3.9 (3.8) |
| Wilson B-factor (Å$^2$) | 44.0 | 87.4 |
| **Refinement** | | |
| Resolution (Å) | 47.0-2.40 | 48.4-2.70 |
| No. reflections | 20408 (2019) | 13994 (1389) |
| $R_{work}$ | 0.221 (0.344) | 0.255 (0.399) |
| $R_{free}$ | 0.267 (0.402) | 0.296 (0.425) |
| No. atoms | | |
| Protein | 3750 | 3605 |
| Peptides | 38 (--PR) | 16 (-EPL) |
| Ligands¤ | 2 | 0 |
| Water | 27 | 6 |
| Clashscore | 2.8 | 7.0 |
| *B*-factors (Å$^2$) | | |
| Protein | 60.5 | 132.3 |
| Peptides | 80.6 | 122.0 |
| Ligand/ion | 36.9 | - |
| Water | 41.1 | 99.3 |
| R.m.s. deviations | | |
| Bond lengths (Å) | 0.008 | 0.004 |
| Bond angles (°) | 0.73 | 0.61 |
| PDB code | 8CBI | 8CBP |

**Table S2. Map correlation coefficients (CC) from polder map calculations of peptide binding to PrfA.**

| Peptide | Modeled peptide in PrfA | CC(1,2) | CC(1,3) | CC(2,3) |
|---|---|---|---|---|
| **LLL** | **LLL (chain C)** | 0.74 | 0.85 | 0.73 |
| **LLL** | **LLL (chain D)** | 0.70 | 0.88 | 0.76 |
| **EVF** | **EVF (chain C)** | 0.55 | 0.87 | 0.62 |
| **EVF** | **EVF (chain D)** | 0.60 | 0.82 | 0.63 |
| **EVFL** | **EVFL (chain C)** | 0.62 | 0.90 | 0.62 |
| **EVFL** | **EVFL (chain D)** | 0.63 | 0.87 | 0.67 |
| **STLL** | **-TLL (chain C)** | 0.53 | 0.82 | 0.58 |
| **STLL** | **-TLL (chain D)** | 0.53 | 0.84 | 0.61 |
| **RGLL** | **RGLL (chain C)** | 0.54 | 0.88 | 0.44 |
| **RGLL** | **RGLL (chain D)** | 0.70 | 0.80 | 0.61 |
| **TKPR** | **--PR (chain C)** | 0.78 | 0.87 | 0.78 |
| **TKPR** | **--PR (chain D)** | 0.73 | 0.80 | 0.75 |
| **TEPL** | **-EPL (chain C)** | 0.57 | 0.83 | 0.68 |
| **TEPL** | **-EPL (chain C in monB)*** | 0.64 | 0.60 | 0.79 |

The CC values help to judge if the density in the polder omit map resembles the included model of the ligand or the bulk solvent. If the CC(1,3) value is > 0.75 and larger than both CC(1,2) and CC(2,3), the density map most likely represents the modelled ligand[1]. The CC values of the polder maps refers to the positions of the final modeled peptide residues. Modeled peptide chains C and D are modeled in PrfA monomers A and B, respectively.
* As a control, the tripeptide modeled in monomer A of PrfA-TEPL was superimposed into monomer B, even though there was no clear density for the ligand in monomer B. The CC values from polder map support our conclusion that no TEPL peptide bound in monomer B.

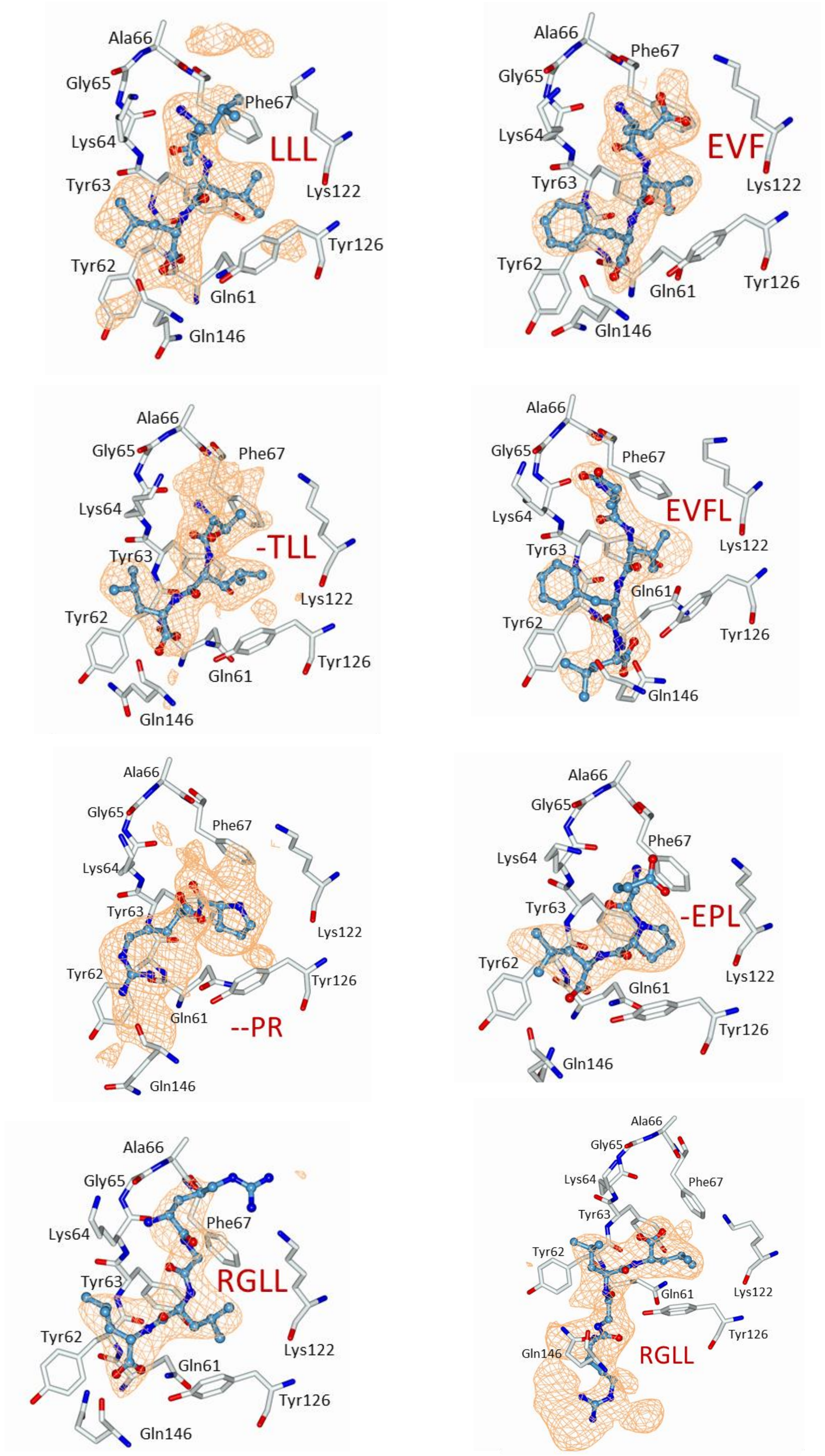

**Fig. S1. Polder maps over the tunnel site in monomer A for PrfA-peptide complexes**. The polder electron density maps are colored in orange and contoured at three times the root-mean-square value of the map. The maps are covering 5 Å around the respective peptides. For the tetrapeptides STLL, TKPR, and TEPL, only the residues 2-TLL-4, 3-PR-4 and 2-EPL-4 could be modelled and refined.

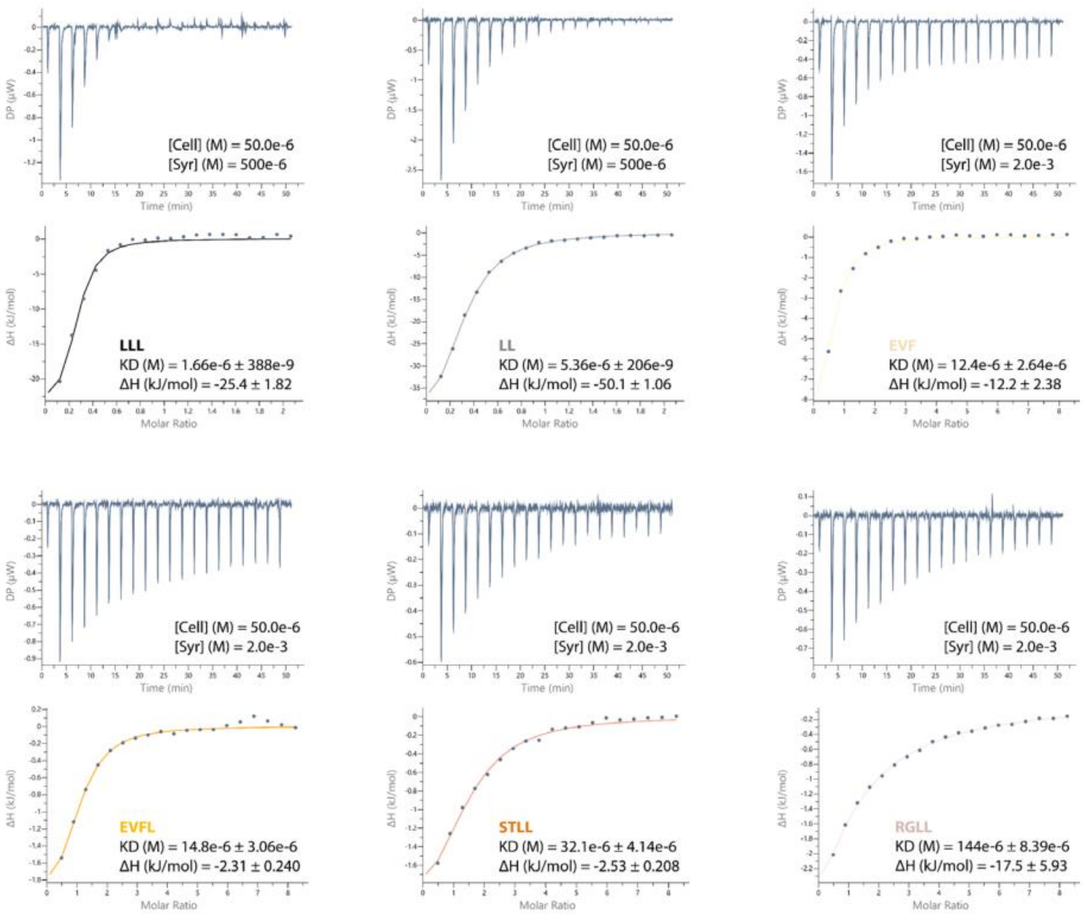

**Fig. S2**. ITC analysis of PrfA-oligopeptide binding interactions. The panels show representative ITC thermograms and corresponding fitted curves generated for the injection of oligopeptides into PrfA solutions. The top panel shows the calorimetric titration, and the bottom panel shows the derived binding isotherm plotted against the titrant molar ratio, with the solid lines representing the best fit to the data.

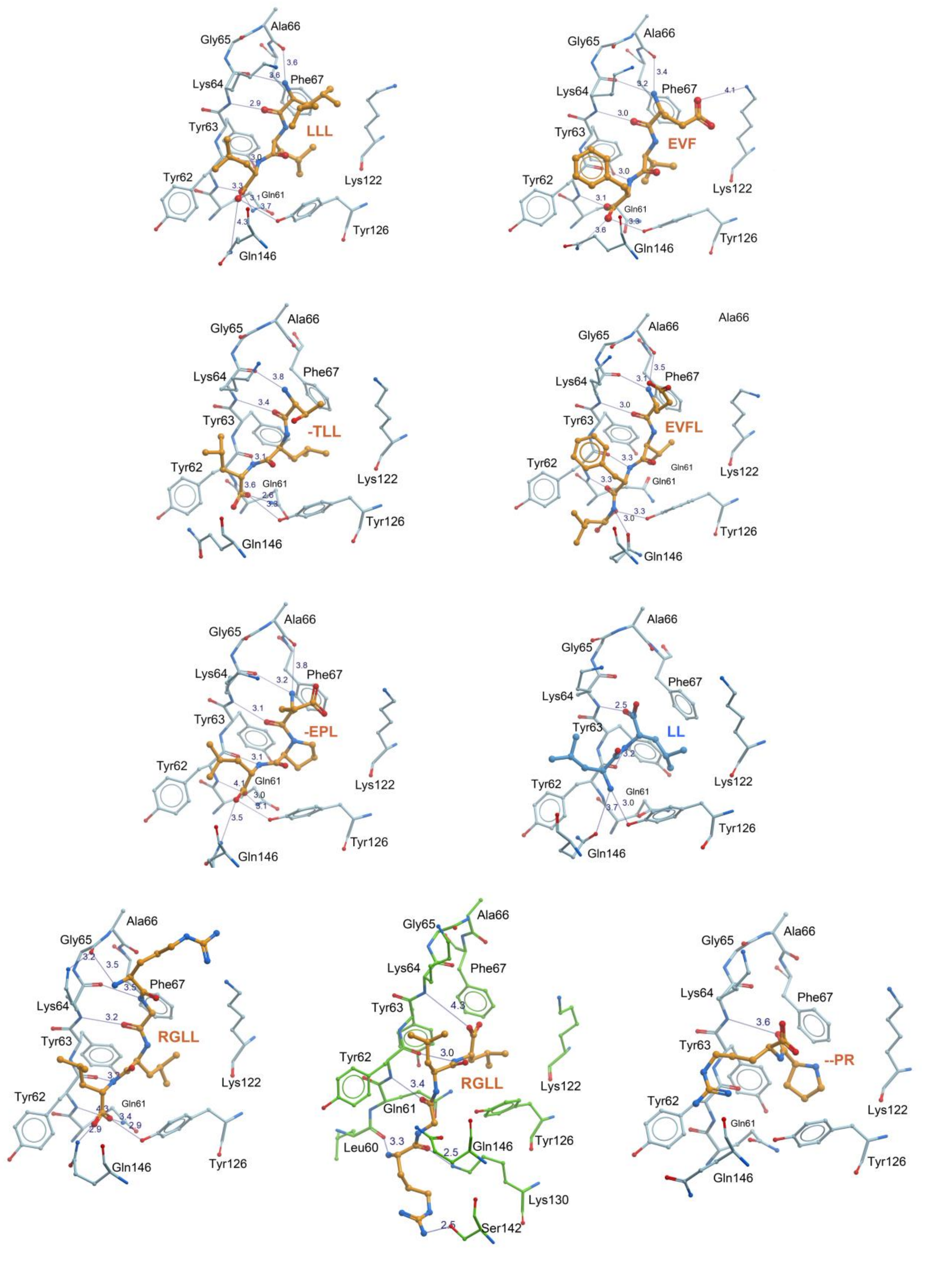

**Fig. S3. Structure of PrfA-ligand complexes.** Binding interactions between peptides and PrfA protein in monomer A. Hydrogen bonds are indicated with thin black lines; bond distances are given in angstrom. The coordinates for the PrfA-LL structure is taken from PDB code 6hck[2]. Binding of RGLL in monomer B is highlighted with green bonds for the protein residues. Only the residues 2-TLL-4, 3-PR-4 and 2-EPL-4 could be modelled and refined for the tetrapeptides STLL, TKPR, and TEPL.

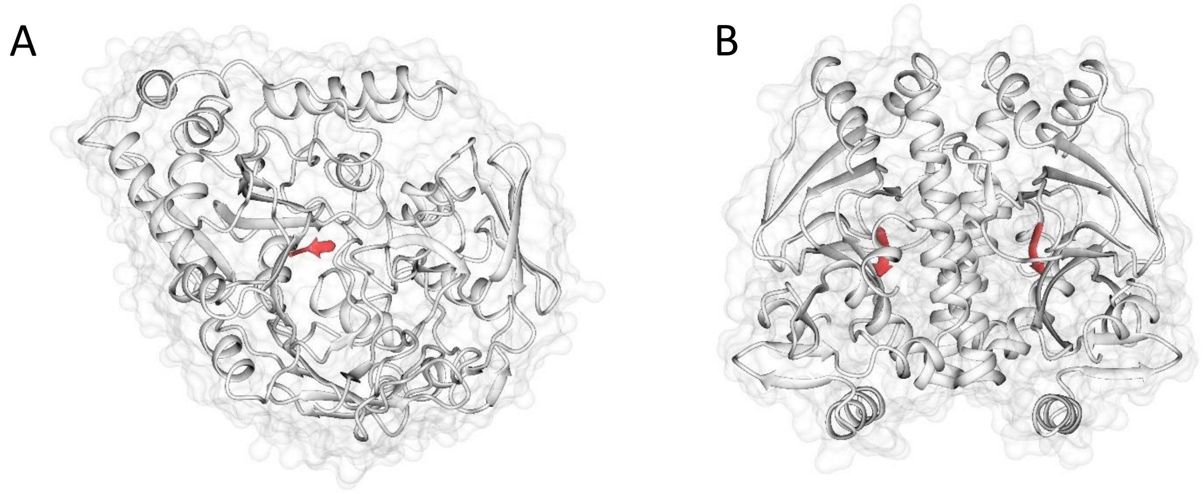

**Fig. S4. Ribbon drawings of peptide conformations in OppA and PrfA.** (A) The oligopeptide binding protein receptor OppA (PDB code 2olb[3]), and (B) PrfA in complex with LeuLeuLeu (this work, PDB code 8cb4), both bind to short peptides in β-strand conformations. The peptides are highlighted in red.

**References:**

1- Liebschner, D. et al. Polder maps: improving OMIT maps by excluding bulk solvent. Acta Crystallogr D Struct Biol 73, 148-157 (2017).
2- Krypotou, E. et al. Control of Bacterial Virulence through the Peptide Signature of the Habitat. *Cell Rep* **26**, 1815-1827 e5 (2019).
3- Tame, J.R., Dodson, E.J., Murshudov, G., Higgins, C.F. & Wilkinson, A.J. The crystal structures of the oligopeptide-binding protein OppA complexed with tripeptide and tetrapeptide ligands. *Structure* **3**, 1395-406 (1995).
